# Smad4 Loss in the Mouse Intestinal Epithelium Alleviates the Pathological Fibrotic Response to Injury in the Colon

**DOI:** 10.1101/2024.03.08.578000

**Authors:** Zahra Hashemi, Thompson Hui, Alex Wu, Dahlia Matouba, Steven Zukowski, Shima Nejati, Crystal Lim, Julianna Bruzzese, Kyle Seabold, Connor Mills, Cindy Lin, Kylee Wrath, Haoyu Wang, Hongjun Wang, Michael P. Verzi, Ansu Perekatt

**Author notes:** **Correspondence:** Ansu Perekatt, Dept. of Chemistry and Chemical Biology, Stevens Institute of Technology, 507 River St., McLean 206, Hoboken, NJ, 07030.

## Abstract

Mucosal healing is associated with better clinical outcomes in patients with inflammatory bowel diseases (IBDs). Unresolved injury and inflammation, on the other hand, increases pathological fibrosis and the predisposition to cancer. Loss of Smad4, a tumor suppressor, is known to increase colitis-associated cancer in mouse models of chronic IBD. Since common biological processes are involved in both injury repair and tumor growth, we sought to investigate the effect of Smad4 loss on the response to epithelial injury. To this end, Smad4 was knocked out specifically in the intestinal epithelium and transcriptomic and morphological changes compared between wild type mice and Smad4 knock out mice after DSS-induced injury. We find that Smad4 loss alleviates pathological fibrosis and enhances mucosal repair. The transcriptomic changes specific to epithelium indicate molecular changes that affect epithelial extracellular matrix (ECM) and promote enhanced mucosal repair. These findings suggest that the biological processes that promote wound healing alleviate the pathological fibrotic response to DSS. Therefore, these mucosal repair processes could be exploited to develop therapies that promote normal wound healing and prevent fibrosis.

**NEW AND NOTEWORTHY:** We show that transcriptomic changes due to Smad4 loss in the colonic epithelium alleviates the pathological fibrotic response to DSS in an IBD mouse model of acute inflammation. Most notably, we find that collagen deposition in the epithelial ECM, as opposed to that in the lamina propria, correlates with epithelial changes that enhance wound healing. This is the first report on a mouse model providing alleviated fibrotic response in a DSS-IBD mouse model *in vivo*.

## INTRODUCTION

Mucosal healing is a prime therapeutic treatment goal for IBD patients^1, 2^. As the largest epithelial lining in the body, the intestinal mucosa provides the critical barrier between the microflora in the intestinal lumen and the gut-associated immune cells^3^. Hence, rapid resealing of the epithelial barrier following epithelial injury is essential to restore homeostasis, and to prevent inflammation^4, 5^.

In the instances where the regenerative capacity of the epithelia is compromised or does not suffice to meet the extent of injury, the exposure of the luminal contents to the underlying mesenchyme triggers an inflammatory response^6^. When unchecked, the inflammatory response can lead to fibrosis, which in turn can lead to strictures, a common complication in Crohn’s Disease (CD)^7^. Chronic inflammation can also increase the predisposition to cancer in Ulcerative Colitis (UC) and CD patients^8^.

The DSS-IBD mouse model is a well-established animal model to study both acute and chronic inflammatory responses in the intestine. DSS administration in drinking water is thought to cause injury by penetrating the epithelial membrane^9^; the severity of the injury and inflammation is dependent on the molecular weight of DSS, the amount administered in the drinking water and the length of the treatment^10^. Restorative response to DSS-induced injury involves several steps: hemostasis to seal breached vasculature, cell cycle arrest, spreading of the epithelial cells to cover the denuded area, and differentiation to restore the epithelial integrity^11^. In parallel, the immune cells are activated to clear the infiltrated immunogens from the injury^12^. However, an unresolved injury due to compromised regenerative capacity can cause fibrosis, a pathological wound-healing response wherein fibroblasts and collagen replace the epithelium^13^.

Chronic injury also increases the predisposition to cancer. The wound healing process is known to causes systemic changes that desensitize the immune cells, thereby creating an immunosuppressive milieu that permits the growth of neoplastic cells^14, 15^. Smad4 is a tumor suppressor^16–19^, and loss-of-function mutation(s) in Smad4 is one of the major drivers of colon cancers^20^. While Smad4 loss alone in the intestinal epithelium does not affect the gross phenotype^21, 22^, it increases colitis associated cancer^23, 24^.

To distinguish the initial DSS-induced injury response from the long-term tumorigenic effect of Smad4 loss in the intestinal epithelium, we used a DSS-IBD mouse model of acute inflammation. Smad4 was knocked out specifically in the intestinal epithelium (Smad4^IEC-KO^) followed by DSS treatment. DSS-induced transcriptomic changes specific to the epithelium were evaluated after three days of treatment, and the molecular and gross phenotypic changes were evaluated after three and seven days of DSS treatment.

We find that Smad4 loss specific to the intestinal epithelium alleviates the fibrotic response to DSS in the mouse colon. Our findings reveal the molecular changes that alter the epithelial extracellular matrix (ECM) in the epithelium of the Smad4^IEC-KO^ mice. We also provide evidence for improved mucosal healing that can be ascribed to the alleviated pathological response to DSS in the Smad4^IEC-KO^ mouse colon.

## MATERIALS AND METHODS

### Materials

Antibodies and reagents are summarized in the supplementary tables 1-3.

### Animals and Experimental Protocol for Inducing Colitis

The animal experiments were conducted according to the protocol approved by the IACUC of Stevens Institute of Technology and Rutgers. All mice were kept under a 12-h light/dark cycle and harvested around mid-day to prevent diurnal variations. To create the Smad4 knockout conditional mutant, the *Villin-Cre^ERT^*^2^ transgene^25^ was integrated into Smad4^f/f^^26^ mice conditional-mutants and controls. To induce epithelial-specific loss of Smad4, 0.05 g/kg tamoxifen mice per day was injected intraperitonially for four consecutive days. 2.5% DSS (40 kDa) was administered in drinking water for *ad libitum* drinking for three or seven consecutive days, depending on the purpose of the experiment. The treatment regimen for Smad4 deletion and DSS treatment are shown (Figure S1A). The animals used were gender and age matched across the different treatments. All animals used were between eight and fourteen weeks of age.

### Colon Epithelial Isolation for RNA Extraction and Sequencing

Freshly harvested colon was flushed with PBS, filleted open, cut into ∼ 1 cm pieces, and the epithelia separated from the underlying mesenchyme using EDTA chelation as follows: the intestinal pieces were incubated with 5 mM/L EDTA/PBS for 50 minutes at 4°C, and then shaken to separate the epithelium from the underlying mesenchyme. The colonic crypt epithelium was isolated after filtering through a 70-micron filter. The isolated crypts were washed twice with ice-cold PBS by spinning at 200 rcf 4°C for two minutes. Any remaining PBS was aspirated prior to solubilizing in TRIzol and RNA extracted as per the manufacturer’s recommendation.

### Protein Extraction

Whole colonic tissue from the proximal colon was used for protein extraction. After flushing the colon with cold PBS, the proximal colon was minced into 1 cm pieces and flash-frozen in liquid Nitrogen and macerated using a pestle and mortar. The macerated tissue was lysed in RIPA buffer (20 mM/L HEPES,150 mM/L NaCl, 1 mM/L EGTA, 1% Triton x-100, 1 mM/L EDTA) containing freshly added protease inhibitors (1x PI, 20mM/L NaF, 1 mM/L Na3VO3, 1 mM/L PMSF), rocked at 4°C for 30’, and spun at 4°C to extract the solubilized protein. The protein concentration was determined using the BCA Protein Assay Kit.

### ELISA for C-Reactive Protein (CRP)

Six micrograms of the protein extract were used for the CRP assay using CRP assay kit. The flash-frozen protein extracts, after normalizing the concentration, were used for the CRP assay. Standards were prepared freshly, and the CRP assay carried out as per manufacture’s recommendation.

### Histology, Immunostaining, and Image Acquisition

Freshly isolated colon was flushed with PBS, opened longitudinally, made into Swiss-rolls and fixed overnight at 4°C with 4% paraformaldehyde in PBS. The fixed tissues were dehydrated in alcohol series and processed in xylene and paraffin prior to embedding them in paraffin. 5-micron sections of the paraffin embedded tissue were used for histological evaluation and immunostaining as previously described^19^. Summary of the antibodies and dilution used are in summarized in the supplementary tables 1-3. Hematoxylin or Methyl green was used as a nuclear counter stain. Brightfield images were obtained using Nikon microscope (Model Eclipse Ci-L; cat. No. M568E) and Nikon camera (DS-Fi3; cat. No. 117837). Fluorescent images were obtained using Zeiss confocal microscope (Carl Zeiss Microscopy GmbH LSM 880). The same laser power and gain was maintained when obtaining fluorescent images. Low-magnification (4X) fluorescent images were obtained using Bio Tek Lionheart FX microscope and camera. Contrast and brightness, when adjusted for bright-field images, were uniformly applied across the treatments being compared.

### BrdU-EdU Pulse Chase and Fluorescent Detection

Mice were injected with 1 mg of EdU and 1 mg of BrdU at one and six hours respectively, prior to sacrificing the mice. Fluorescent immunohistochemistry was used to detect the BrdU-incorporated cells; the EdU-incorporated cells within the same tissue were co-detected with red fluorescent marker as per the manufacture’s recommendation. The distance of migration was calculated by measuring the distance between the top BrdU-positive and the EdU-positive cell relative to the height of the crypt being accounted. The quantification was performed on the images captured from the distal colon. At least 30 open crypts were accounted for each sample.

### Picrosirius Red Attaining and Assessment Of Collagen Proportionate Area

Five-micron sections of paraffin-embedded colons were hydrated and stained for collagen using Sirius red/Fast green kit as per manufacturer’s recommendation. Collagen deposition was assessed only in the mucosae, i.e., collagen in the submucosa, mucosa muscularis and serosa were not accounted. Collagen positive pixels were quantified as described earlier^27^. Briefly, five 10x images from distal colon were randomly selected for the assessment. The images were then deconvoluted to visualize only the collagen positive pixels and quantified to obtain collagen positive pixels per unit area of the mucosa.

### Quantification of Intact Crypts, Crypt-Less Mucosal Regions

Ten randomly selected 20x images of α-SMA-immunostaining the distal colon was used. The number of intact crypts per unit area only in the mucosa were counted manually. The same images were used to quantify the crypt-less regions per unit area of the mucosa, containing α-SMA-positive and α-SMA-negative cells.

### Quantification of Col1a1 (Type I Collagen) in the Peri-Crypt Epithelium

Ten 20x images of the distal colon immunostained for Col1a1 for each replicate were used. Col1a1 immunoreactivity in the peri-crypt epithelium of at least fifty open crypts for each replicate were quantified as follows (Figure S3C): A rectangle was drawn around the peri-crypt epithelium to assign the the total area accounted for each crypt. Using ImageJ 1.53t and Zen 3.1 software, the Col1a1 immunoreactive region in the peri-crypt epithelium was outlined. The outlined Col1a1-positive area per unit area of the rectangle was quantified.

### Quantification of BrdU-Labeled Cells to Assess Proliferation

Mice were injected with 1 mg BrdU two hours prior to sacrifice. The BrdU-incorporated cells were detected by BrdU immunohistochemistry. BrdU-positive cells in at least 40 open and intact crypts per replicate of the distal colon were enumerated.

### Statistical Analysis

Statistical analysis was performed using GraphPad Prism 8.0.2. T tests were used to compare the means of data between two groups. Statistical significance was set at P ≤ 0.05.

### RNA-Seq Library Preparation and Sequencing

Quality control, library preparation, and sequencing for the study were conducted by Novogene Corporation Inc. (Sacramento, CA). RNA integrity was initially assessed using the RNA Nano 6000 Assay Kit of the Bioanalyzer 2100 system (Agilent Technologies, CA, USA), and only RNA samples with an RIN value above 4 were selected for further processing. This process involved the purification of mRNA and subsequent cDNA synthesis. After adapter ligation, the cDNA underwent another purification step. The prepared library’s quality was then evaluated using the Agilent Bioanalyzer 2100 system. Finally, sequencing was performed on an Illumina Novaseq platform, generating 150 bp paired-end reads. The raw sequencing reads, provided in FASTQ format, were processed by Novogene using their in-house Perl scripts to obtain clean reads. These clean reads were then aligned to the mm9 mouse reference genome using Hisat2 (v2.0.5)^28^. Subsequently, the aligned reads were quantified for gene expression levels using featureCounts (v1.5.0-p3)^29^.

### Bioinformatics

Differential expression analysis for pairwise comparisons was performed using the DESeq2 (v1.40.2) package^30^ in R (v4.3.1)^31^. P-values were adjusted using the Benjamini-Hochberg method to control the false discovery rate^32^. Genes with an adjusted p-value (padj.) of less than 0.05 were considered significantly differentially expressed.

Gene Ontology (GO)^33^ enrichment analysis was performed on the significantly differentially expressed genes (DEGs) using the clusterProfiler (v4.9.4) R package^34^. P-values obtained from the hypergeometric test were adjusted using the Benjamini-Hochberg method for multiple hypothesis testing. GO terms with an adjusted p-value (padj) less than 0.05 were deemed significantly enriched.

Normalized counts for Gene Set Enrichment Analysis (GSEA) were obtained using the counts (dds, normalized=TRUE) function from DESeq2, and GSEA (v4.3.2) was conducted using the Broad Desktop Application^35, 36^. Mouse Ensembl IDs were mapped to Human Gene Symbols, and genes were ranked using the Signal2Noise metric. The analysis utilized the weighted enrichment statistic and 10,000 gene set permutations, with gene sets filtered by size (minimum 15, maximum 500). Gene sets with a false discovery rate (FDR) q-value below 0.05 were identified as significantly enriched.

Gene sets were sourced from the Molecular Signatures Database (MSigDB)^37^. Heatmaps of core-enriched genes and UpSet plots were generated using the ComplexHeatmap package in R^38^. UpSet plots were used to determine the uniquely enriched gene sets across each pairwise comparison that was performed.

Ingenuity Pathway Analysis (IPA)^39^ was performed with the differentially expressed genes (DEGs) identified in our study. DEGs with |Log2FoldChange| ≥ 0.584 were uploaded to IPA for functional annotation and regulatory network analysis.

## RESULTS

### Epithelial Loss of Smad4 In The Intestine Alleviates The Pathological Response To DSS

To determine the effect of epithelial-specific loss of Smad4 on DSS treatment, we knocked out Smad4 only in the epithelium using the Villin promoter-driven, tamoxifen inducible cre-recombinase^25^. Four days of Tamoxifen injection caused epithelial specific loss of Smad4 loss throughout the intestinal epithelium(S1B). The Smad4 knocked out (Smad4^IEC-KO^) and the wildtype (WT) mice were then treated with 2.5% DSS (40 kDa) for a week and the phenotypic and histological changes were evaluated. Bloody stools and loss in body weight were apparent within 7 days. However, the loss in the body weight in response to DSS treatment was significantly lower in the Smad4^IEC-KO^ mice as compared to that in the wild type mice (Figure 1A). The severity of colitis response to DSS, assessed by colon length and histology, was also reduced in the Smad4^IEC-KO^ mice (Figures 1B & D). Furthermore, the number of intact epithelial crypts remaining after a week of DSS treatment was higher in the Smad4^IEC-KO^ colon (Figure 1C). These observations collectively indicate that Smad4 loss in the intestinal epithelium has a protective effect against DSS-induced damage.

**Figure 1.**
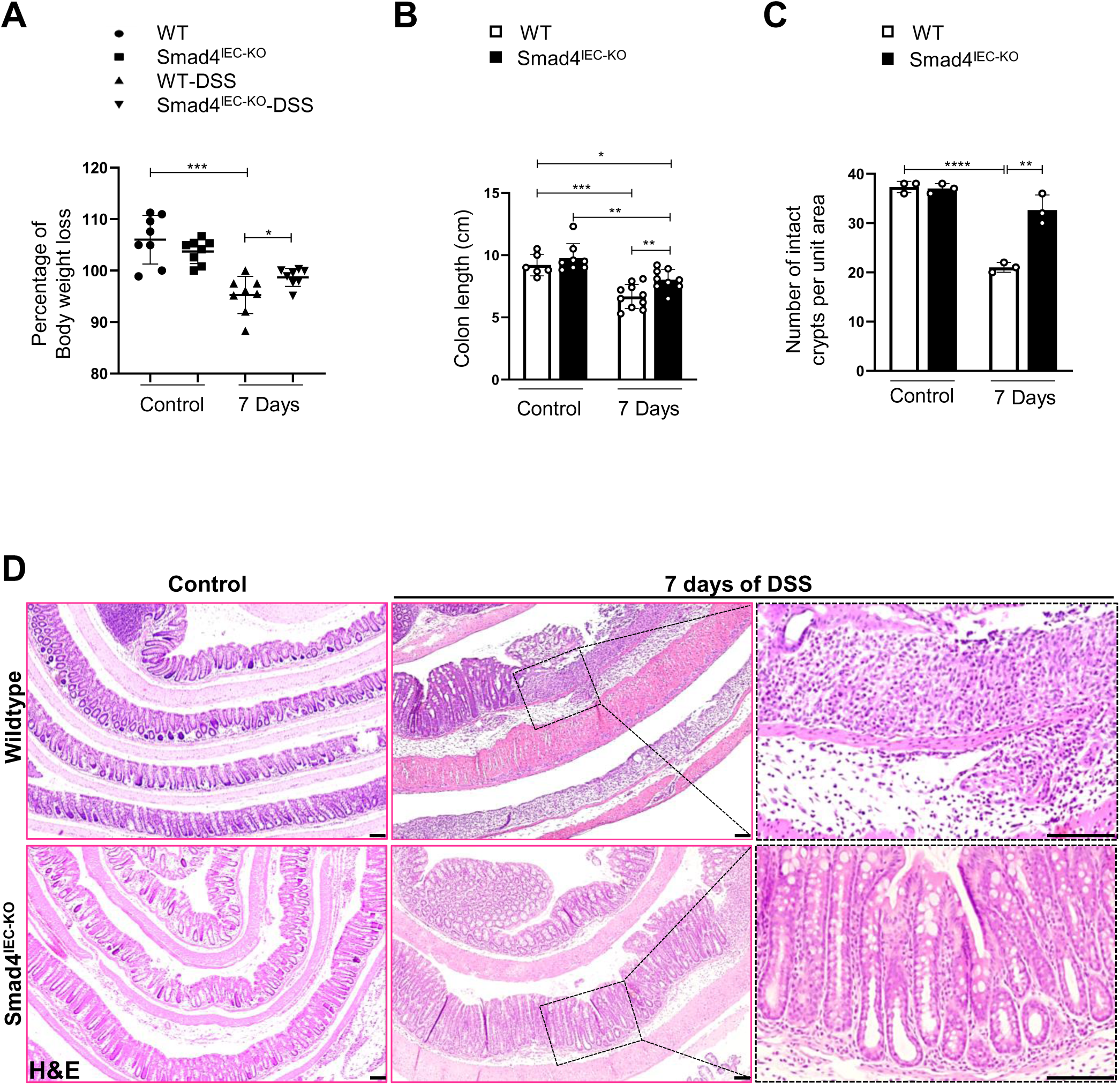
Attenuated pathological response to seven days of DSS in the Smad4^IEC-KO^ colon: DSS-induced (A) weight loss (n=35: 3 replicates per WT (wild type), 3 replicates per Smad4^IEC-KO^, 12 replicates per WT-DSS-7 days, and 17 replicates per Smad4^IEC-KO^-DSS 7 days) The legends in the jitter plot indicate the body weight. (B) colon length (n=33: 6 replicates per WT, 8 replicates per Smad4^IEC-KO^, 10 replicates per WT-DSS 7 days, 9 replicates per Smad4^IEC-KO^-DSS 7 days). (C) Number of intact crypts per unit area DSS treatment. (D) H&E staining showing the histology of the DSS-treated colon (n=12: 3 replicates per treatment; *,**,*** and **** denote p < 0.05, p < 0.01, p < 0.001 and p <0.0001, respectively). Scale bars, 100 μm.

### Reduced Fibrotic Response to DSS Treatment in The Smad4^IEC-KO^ Colon

Fibrosis is a pathological wound healing response in which the damaged epithelial tissue is replaced by fibrotic tissue^40, 41^. Since the Smad4^IEC-KO^ colon showed higher retention of intact crypts after DSS treatment, we assessed the fibrotic response. Fibrosis is characterized by an increase in mesenchymal cells, including myofibroblasts, and deposition of ECM components^42, 43^. Hence, we first assessed collagen deposition, a hallmark of fibrosis^44^ using Picrosirius red (PSR) staining (Figure 2A). Digital quantification of the PSR-stained region of the mucosa revealed a significant increase of the collagen proportionate area (CPA)^27^ after DSS treatment in the wildtype, but not in the Smad4^IEC-KO^ colon mucosa (Figure 2B). Next, we tested for fibroblasts, a major source of collagen^45^ by immunostaining for α-SMA^46, 47^. Immunoreactivity for α-SMA in the wildtype colon mucosa was restricted to the region lacking intact crypts (Figure 2C), which was higher in the DSS-treated wildtype colon (Figure 2D). In the Smad4^IEC-KO^ colon mucosa, however, α-SMA immunoreactivity was detected in the peri-crypt epithelium (Figure 2C), suggesting activation of sub-epithelial myofibroblasts^46, 48^ in the peri-crypt epithelium of the Smad4^IEC-KO^, but not the wildtype colon.

**Figure 2.**
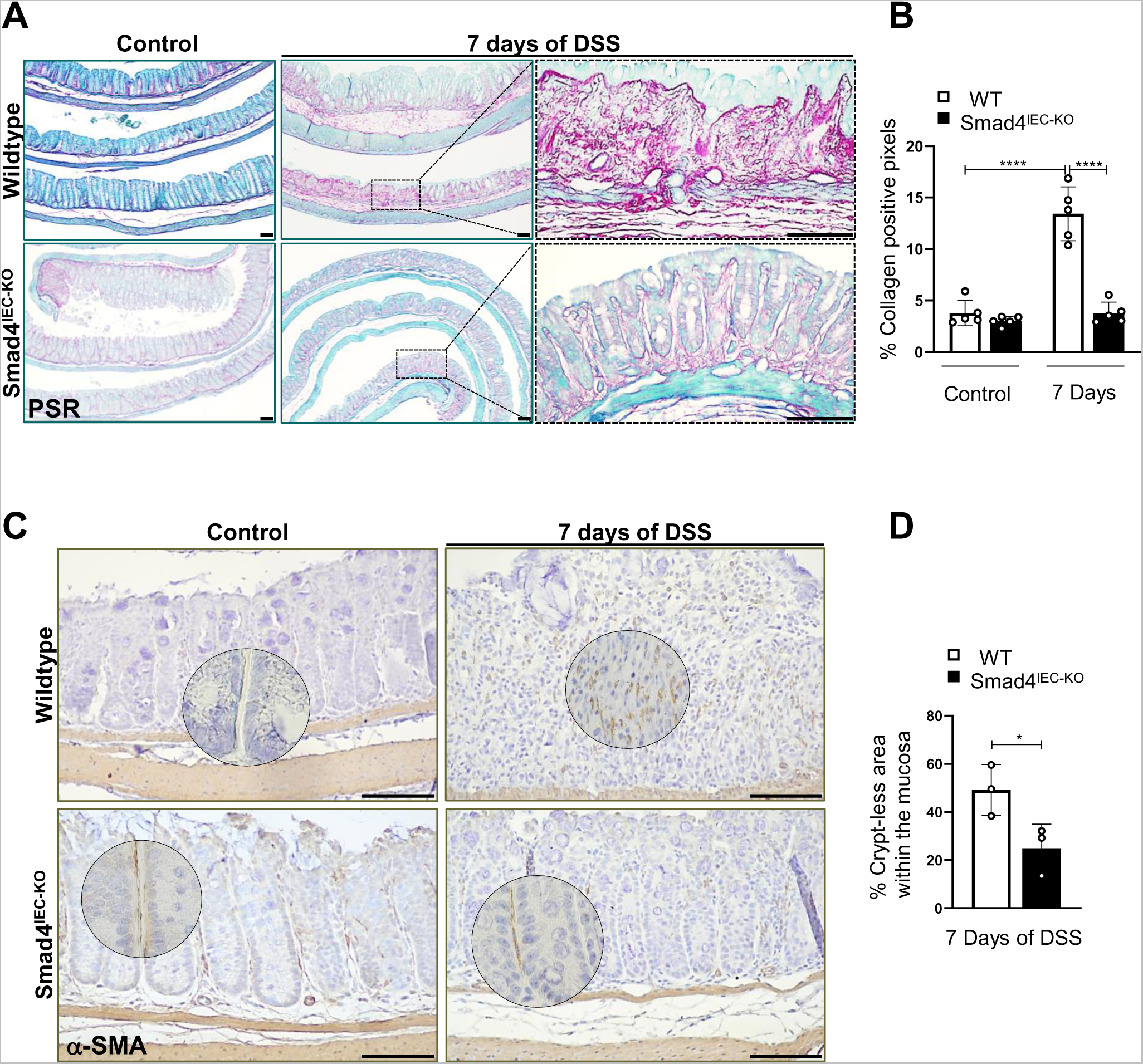
Dampened fibrotic response to DSS in the Smad4^IEC-KO^ colon: (A) Picrosirius red (PSR) staining showing collagen deposition, and (B) quantification of the collagen deposition, assessed by collagen proportionate area (CPA). (C) α-SMA immunoreactivity indicating activated fibroblasts. Scale bars, 100 μm. (The magnified regions are encircled), and (D) the relative proportion of the α-Sma-positive areas in the mucosal regions devoid of intact crypts (n=12: 3 replicates per treatment; * and **** denote p < 0.05 and p <0.0001, respectively). Scale bars, 100 μm.

### The Proliferative Zone of the Smad4^IEC-KO^ Colon Epithelium Expands after DSS Treatment

DSS is known to breach the epithelial surface, thereby exposing the luminal contents of the colon to the subepithelial surface, ultimately triggering inflammation and colitis^49^. Hence, faster resealing of the breached epithelium can minimize the DSS-induced inflammation and colitis. To investigate the wound healing responses toward resealing of the epithelium, we first assessed epithelial proliferation after DSS. No significant difference between the wild type and Smad4^IEC-KO^ colon epithelium was observed after three days of DSS treatment (Figure 3A). After seven days of DSS treatment, however, the number of BrdU-positive cells decreased in the intact crypt epithelium of the wild type mice (Figure 3B). No such decrease was observed in the Smad4 ^IEC-KO^ colon epithelium after seven days of the DSS treatment, indicating that proliferation was uninterrupted by the DSS treatment in the Smad4^IEC-KO^colon epithelia. Consistent with this, the GO-term “Cell Cycle Arrest” was negatively enriched, while Myc target gene signatures were positively enriched in the DSS-treated Smad4^IEC-KO^ colon epithelium compared to its wildtype counterpart (Figures S2A & C). Furthermore, the transcript levels of *Cdkn1a*, encoding the cell cycle inhibitor p21, are significantly lower in the Smad4^IEC-KO^ colon epithelium compared to the wildtype (Figure S2B). This finding further supports the notion that proliferation in the Smad4^IEC-KO^ colon epithelium is uninterrupted by DSS treatment, in contrast to the reduction in proliferation observed after DSS in the wildtype epithelium^11, 50^(Figure 3B).

**Figure 3.**
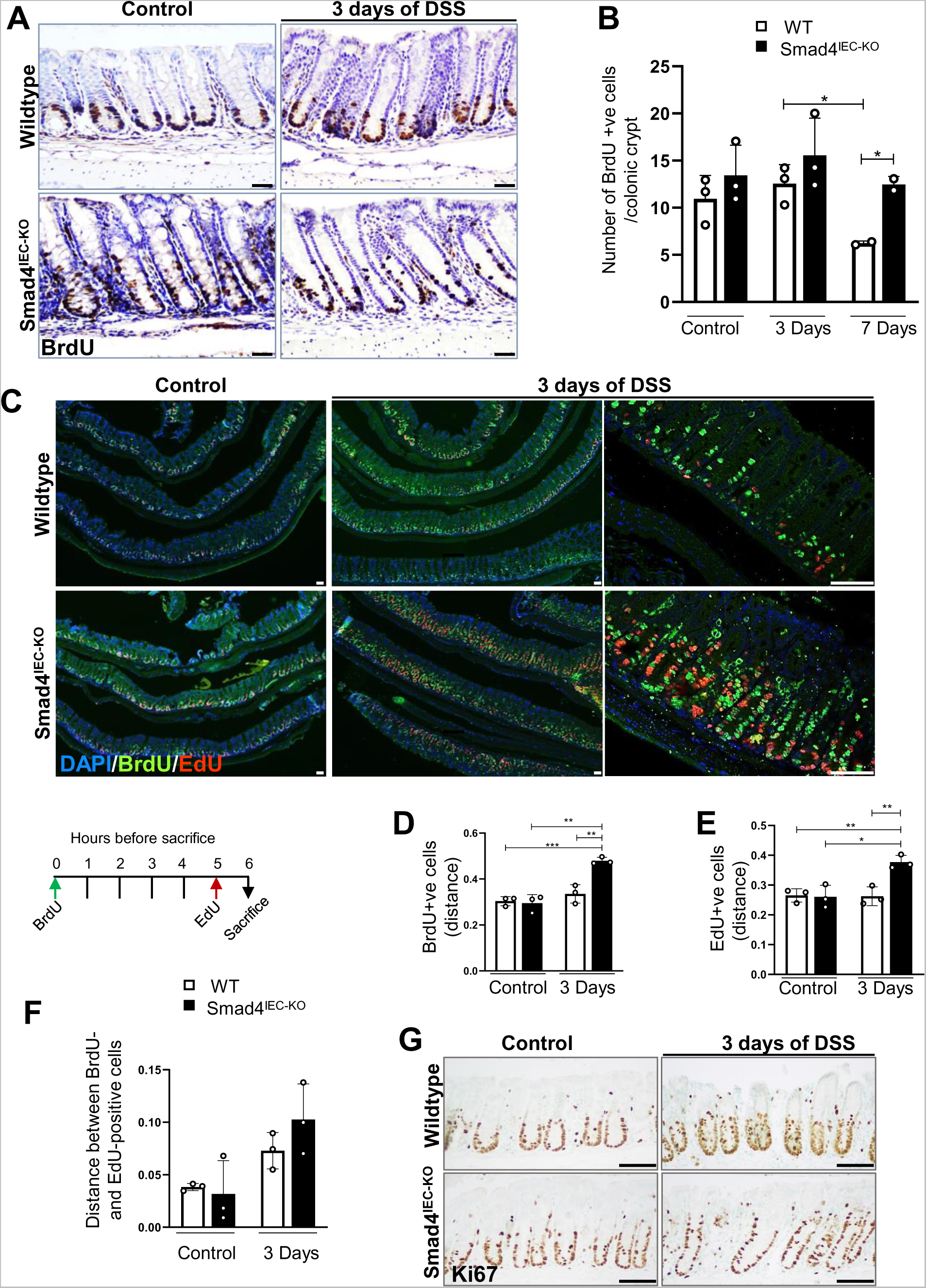
Expansion of the proliferative zone in the Smad4^IEC-KO^ colon epithelium upon three days of DSS treatment. (A) BrdU IHC showing proliferating crypt epithelial cells in S phase pulse-chased for two hours, and (B) their quantification (n=16: 3 replicates per all treatments except the for the 7-day timepoint for which is 2 replicates per treatment (C) BrdU and/or EdU incorporated cells in the crypt epithelium after pulse-chase with BrdU and EdU for six- and one-hours, respectively; below is the schematic showing the timeline of BrdU and EdU pulse-chase. (D) Relative height at which the BrdU-labeled cells (green) are found in the crypts after 6 hours of BrdU pulse-chase. (E) EdU labeled cells (red) are found in the crypts after one hour of Edu pulse-chase. (F) migration assessed by the distance between the BrdU- and the Edu-positive zones within the same crypt relative to the crypt height. (G) Representative images of Ki67 IHC showing proliferative zone within the crypts. (n=12: 3 replicates per treatment; (X-axis on D, E, and F represents the ratio of the distance relative to the crypt height). * and **, *** denote p < 0.05, p < 0.01 and p < 0.001, respectively). Scale bars, 50 μm.

Since the BrdU-labeled cells were detected higher up in the crypts of Smad4^IEC- KO^colon, we tested for increased migration after DSS treatment. To this end, double pulse labeling with EdU and BrdU was performed at one and six hours prior to the sacrifice, respectively (Figure 3C). Both, BrdU and EdU positive cells were detected significantly higher up in the DSS-treated Smad4^IEC-KO^ colon epithelium (Figures 3D & E). However, the distance between the BrdU- and EdU-labeled cells in the Smad4^IEC-KO^ compared to its wildtype counterpart, was not significant (Figure 3F). These observations indicate an expansion of the proliferative zone within the Smad4^IEC-KO^ crypt epithelium after DSS treatment. Consistent with this, the proliferative marker, Ki67 was also detected higher up in the crypts (Figure 3G).

Collectively, these data indicate expansion of the proliferative zone in the crypt epithelium of the Smad4^IEC-KO^ colon after DSS treatment without affecting migration significantly.

### The Crypt Epithelia of The Smad4^IEC-KO^ Colon Display ECM Alterations that Support Wound Healing Response

Next, we investigated the transcriptomic changes specific to the epithelium that can be attributed to the alleviated pathological response to DSS in the Smad4^IEC-KO^ colon.

To this end, we performed mRNA-sequencing analysis on the colon epithelia from the wildtype and Smad4^IEC-KO^ mice after three days of DSS treatment – a time point at which DSS-induced molecular changes can be detected without gross morphological changes or epithelial loss^49, 51^. Over-representation analysis (ORA) on the differentially expressed genes (DEGs) showed Gene Ontology (GO) terms associated with ECM organization, collagen components, and wound healing to be amongst the most enriched GO terms in the DSS treated Smad4^IEC-KO^ epithelia compared to its wildtype counterpart (Figure 4A). Consistent with this, the DSS treated Smad4^IEC-KO^ epithelium showed enrichment for molecular signatures of ECM constituents (Figure 4B), and higher transcript levels of genes encoding various collagens (Figure 4E). Furthermore, stronger Immunoreactivity for type I collagen was detected in the peri-crypt epithelium of the DSS-treated Smad4^IEC-KO^ colon (Figures 4C & D), suggesting increased collagen in the crypt epithelial ECM of the Smad4^IEC-KO^ colon.

**Figure 4.**
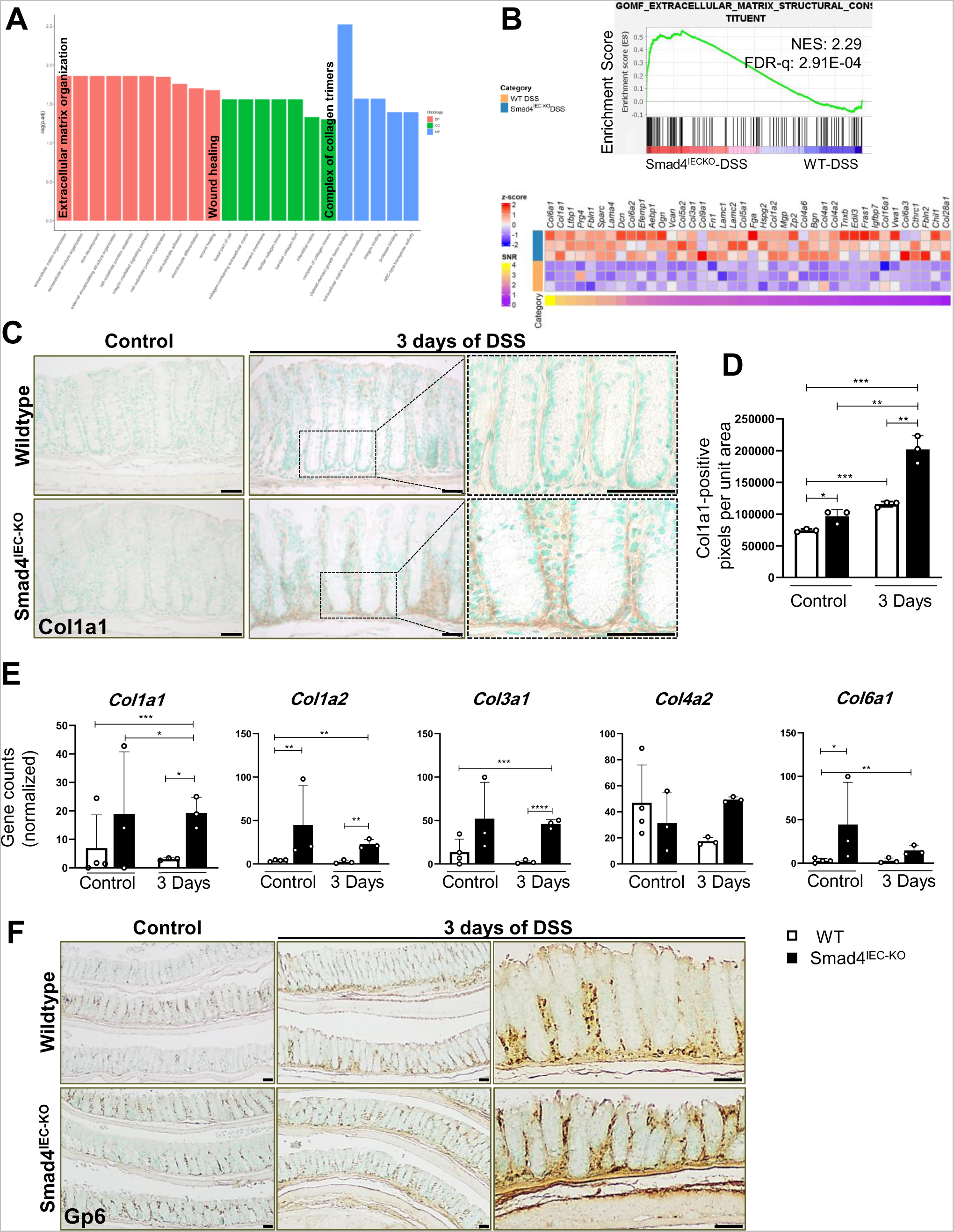
ECM alterations in the DSS treated Smad4^IEC-KO^ colon supporting wound healing response. (A) Top GO terms enriched in the epithelial transcriptome of DSS-treated Smad4^IEC-KO^ colon compared to its wild-type counterpart. (B) GSEA plot showing signatures of ECM organization in the Smad4^IEC-KO^ colon epithelia; the relative levels of the core- enriched genes are visualized in the heatmap. (C) Representative images of IHC for Type I Collagen and, (D) quantification of the immunoreactive pixels in the epithelial ECM. (E) Increased transcript levels of the genes encoding various collagens: *Col1a1*, *Col1a2*, *Col3a1*, *Col4a2* and *Col6a1* in the Smad4^IEC-KO^ colon epithelia. (n=13: 4 replicates per WT and 3 replicates per the rest of the treatments); p.adj. values obtained from DEG analysis were used to determine the significance (F) Gp6 immunoreactivity in the colon before and after three days of DSS. (n=12: 3 replicates per treatment; *,**,*** and **** denote p < 0.05, p < 0.01, p < 0.001 and p <0.0001, respectively). Scale bars, 50 μm. NES = normalized enrichment score, FDR- q = False discovery rate q-value.

To investigate the functional implication of the ECM-related gene signatures specifically in the epithelium, we performed Integrated Pathway Analysis (IPA) on the DEGs in the DSS-treated Smad4^IEC-KO^ colon epithelium versus its wild-type counterpart. Collagen-mediated Gp6 signaling was amongst the most enriched signaling pathways in the DSS-treated Smad4^IEC-KO^ colon (Figure S3A & B). Collagen is a ligand for the Gp6 receptor on platelets. Activation of Gp6 signaling following engagement of Gp6 receptor with collagen causes platelet aggregation^52^ to seal vascular breaches and thus, promote wound healing. While no differences in Gp6 immunoreactivity was detected (Figure 4F), higher collagen in the ECM of Smad4^IEC-KO^ epithelia might increase engagement of the Gp6 receptor in the peri-crypt epithelium, contributing to faster resealing of vascular breaches near the epithelium, and thus, enhanced mucosal healing in the Smad4^IEC-KO^ colon (Figure S3D).

### The Smad4^IEC-KO^ Colon Display Dampened Inflammatory Response To DSS

DSS-induced epithelial damage can trigger inflammatory signaling in the neighboring epithelial cells via epithelial damage associated molecular patterns (DAMPs)^53^. Hence, we sought to determine the contribution of the epithelia to the inflammatory response. GSEA showed differences in the gene signatures associated with inflammation after DSS treatment: Signatures of the pro-inflammatory interferon alpha (IFNα) and interferon gamma (IFNγ)^54, 55^ responses were significantly enriched after DSS treatment in the wildtype intestinal epithelium (Figure 5A). When compared to the DSS-treated wildtype epithelium the DSS-treated Smad4^IEC-KO^ colon epithelium showed negative enrichment for the IFNα and IFNγ gene signatures (Figure 5B).

**Figure 5.**
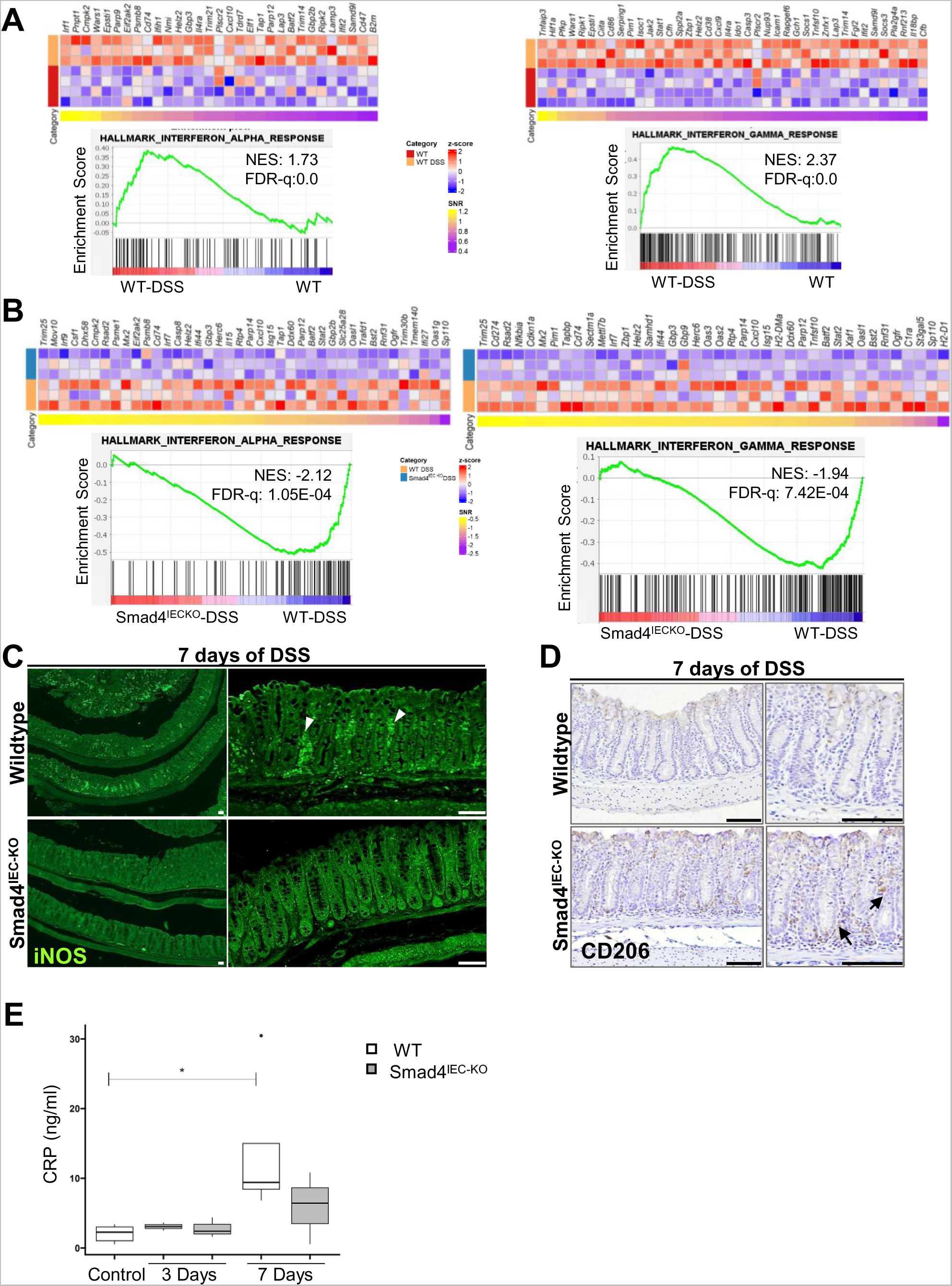
Dampened inflammatory response to DSS in the Smad4^IEC-KO^ colon. (A) GSEA plot showing positive enrichment of IFN-α, and IFN-ψ, signatures after three days of DSS treatment in the wild-type colon epithelium; the corresponding heatmap on the right indicates the core-enriched genes (n=7: 4 replicates per WT and 3 replicates per WT-DSS). (B) GSEA plots showing negative enrichment for IFN-α and IFN-ψ signatures in the Smad4^IEC-KO^ 3-day DSS- treated compared to DSS-treated wild-type colon; the corresponding heatmap above indicates the core-enriched genes (n=6: 3 replicates per treatment). (C) Increased immunoreactivity for iNOS, a pro-inflammatory marker (the right panel shows higher magnification views), and (D) IHC for CD206, an anti-inflammatory M2 macrophage marker, in the DSS- treated wild-type and Smad4^IEC-KO^ colon (black arrows); the right panel shows higher magnification views. (n=6: 3 replicates per treatment). (E) Increased CRP levels in colonic tissues from wild type mice after seven days of DSS treatment (n=18: 5 replicates per WT, 4 replicates per WT- DSS 7 days and 3 replicates per other treatments). (* denotes p < 0.05). NES = normalized enrichment score, FDR q= False discovery rate q-value. Scale bars, 50 μm.

To test if these epithelial changes affect the immune cell infiltration, we probed for pro- and anti-inflammatory markers of the innate immune response. Increased iNOS immunoreactivity was detected in the DSS-treated wild type colon compared to that in the DSS-treated Smad4^IEC-KO^ colon, indicating increased pro-inflammatory response to DSS in the wildtype colon (Figure 5C). Conversely, immunoreactivity for the anti-inflammatory M2 macrophage CD206, was higher in the DSS-treated Smad4^IEC-KO^ colon compared to that in the wildtype colon (Figure 5D), suggesting suppression of pro-inflammatory responses in the Smad4^IEC-KO^ colon after DSS treatment. To determine if the pro-and anti-inflammatory changes correlated with the level of inflammation, we next assayed for C-Reactive Protein (CRP), an indicator of inflammation. DSS treatment increased the CRP levels in the WT colon, but not in the Smad4^IEC-KO^colon (Figure 5E). These data collectively indicate that Smad4 loss in the epithelium dampens the inflammatory response to DSS.

## DISCUSSION

Here, we show that loss of Smad4 in the mouse colonic epithelium alleviates the pathological fibrotic response to DSS. Our findings suggest that uninterrupted proliferation and expansion of the proliferative zone in the Smad4^IEC-KO^ epithelium after DSS contributes to the alleviated pathological response. Additionally, we find transcriptional changes in the Smad4^IEC-KO^ epithelium that alter the epithelial ECM, potentially contributing to enhanced mucosal healing.

Increased collagen in the epithelial ECM of the Smad4^IEC-KO^ colon might contribute to enhanced wound healing response to DSS in two respects: hemostasis and epithelial restoration. Hemostasis is one of the earliest restorative responses that reseals the vascular breach caused by DSS treatment and the consequent neutrophil infiltration^56–58^. Collagen-mediated Gp6 signaling causes platelet aggregation at the site of injury to effect hemostasis^59–61^. Our findings on collagen-mediated Gp6 signaling (Figures S3A, B & S3H) and wound healing (Figures S3D & S3H) therefore suggests that increased collagen in the epithelial ECM (Figures 4C & D) might contribute to better resolution of the vascular damage in the Smad4^IEC-KO^ colon.

ECM could also contribute to the epithelial restoration after damage. Type I collagen in the epithelial ECM is known to promote proliferation^62^ and migration, as well as promote differentiation and homeostasis in the absence of cell-cell contact^63, 64^. Since, the denuded epithelium resembles epithelia lacking cell-cell contact, it is plausible that the increased type I collagen in the Smad4^IEC-KO^ peri-crypt epithelium restores homeostasis faster. The uninterrupted proliferation, and increased expression of E-Cadherin^65^ and Keratin20^66^ expression (Figures S3F & E) in the DSS-treated Smad4^IEC-KO^ colon epithelium suggested repair responses that restore homeostasis. Although no significant increase in migration was observed in the Smad4^IEC-KO^ colon (Figure 3F), the expanded zone of proliferation (Figures 3C-E) is likely to cover the denuded regions in the crypt epithelium, thereby contributing to faster epithelial restoration after damage.

One of the most interesting findings was the altered distribution of α-SMA-positive myofibroblasts. α-SMA-positive myofibroblasts, indicating activation of myofibroblasts,^46, 48^ were more apparent in the peri-crypt epithelial region of the Smad4^IEC-KO^ colon than in the wildtype (Figure 2C). Signaling from epithelia is known to activate sub-epithelial myofibroblasts and cause collagen secretion^67^. Thus, the increased collagen in the peri-crypt epithelial region of the Smad4^IEC-KO^ colon could be attributed to activated subepithelial myofibroblasts. This notion is consistent with the higher transcript levels of *Pdgfa* in the Smad4^IEC-KO^ epithelium (Figure S3G), which encodes the ligand for the *Pdgfra* receptor expressed by fibroblasts^68, 69^.

In the wildtype colon, however, α-SMA-positive fibroblasts were restricted to the mucosal region lacking intact epithelial crypts (Figure 2C). The loss of colonic crypt epithelium in the wildtype thus correlates with increased α-SMA-positive fibroblast-containing mesenchymal regions. Thus, the α-SMA-positive fibroblasts in the crypt-less mesenchymal regions of the wildtype colon could be ascribed to the collagen deposition in the wildtype colon (Figures 2A & B). These distinct observations leads to the speculation that α-SMA-positive fibroblasts in the crypt-less interstitium and the subepithelial myofibroblasts in the peri-crypt epithelial region might have contrasting effects on mucosal healing: Collagen deposition in the crypt-less interstitium by α-SMA- positive fibroblasts promotes pathological fibrosis, whereas collagen deposition in the peri-crypt epithelium by the α-SMA-positive sub-epithelial myofibroblasts promotes restorative mucosal healing. Addressing the fundamental basis for the differential activation of the fibroblasts and the epithelial signals involved in the activation and deposition of the various ECM components is essential to, both prevent pathological fibrosis and promote restorative mucosal healing.

Our studies are distinct from the previously reported tumorigenic effect of Smad4 loss in mouse models of chronic inflammation^23, 24, 70^. Previous studies used haploinsufficiency^71^, or partial deletion^23^, of Smad4 loss in the epithelium and show that Smad4 loss promotes colitis and colitis-associated cancer^23, 24^. Furthermore, in contrast to previous studies that used DSS-IBD mouse models of chronic inflammation, our study employed a model of acute inflammation ^23,24,70^. However, given the role of Smad4 in genomic stability^21, 22, 72^ and tumor suppression^16, 17^, it is not surprising that Smad4 loss led to tumorigenesis in the chronic inflammatory DSS-IBD mouse model, especially in the presence of a DNA-damaging agent such as AOM.^23,24,70^.

Our studies using a DSS-IBD model of acute inflammation (Figure S1A) demonstrate that Smad4 loss in the colon epithelium protects against pathological fibrosis. While enhanced epithelial integrity might have contributed to an alleviated pathological response and dampened inflammatory response, the infiltration of the anti-inflammatory M2 macrophages^73^ in the Smad4^IEC-KO^ colon after DSS (Figure 5C) implicate immunomodulation in suppressing the pathological inflammatory response. This observation, hence, warrants further investigation especially since anti-inflammatory immune infiltration has been implicated in preventing fibrosis^74^.

In summary, our study underscores the importance of the wound healing response in restoring the epithelial integrity after injury, and in preventing the fibrotic response to injury in a DSS-IBD mouse model of acute inflammation. Several biological processes are shared between wound healing and tumor progression^75–77^. Hence, unraveling the biological processes that decouple wound healing from tumor progression will pave the path to therapeutic strategies to promote mucosal healing and prevent pathological fibrosis.

## GLOSSARY

α-SMA: α-Smooth Muscle Actin
AOM: Azoxymethane
BCA: Bicinchoninic acid
BrdU: Bromodeoxyuridine
CD: Crohn’s Disease
CPA: Collagen Proportionate Area
CRP: C-reactive Protein
DEG: Differentially Expressed Genes
DSS: Dextran Sulfate Sodium
ECM: Extracellular Matrix
EDTA: Ethylenediaminetetraacetic acid
EdU: 5-ethynyl-2’-deoxyuridine
EGTA: ethylene glycol-bis (β-aminoethyl ether)-N, N,N′,N′-tetraacetic acid
FDR: False Discovery Rate
GO: Gene Ontology
Gp6: Glycoprotein 6
GSEA: Gene Set Enrichment Analysis
H&E: Hematoxylin and Eosin
IBD: Inflammatory Bowel Disease
IEC: Intestinal Epithelial Cell
IFNα: Interferon Alpha
IFNγ: Interferon Gamma
iNOS: inducible Nitric Oxide Synthase
IPA: Ingenuity Pathways Analysis
KO: Knockout
ORA: Over-Representation Analysis
PBS: Phosphate-Buffered Saline
Pdgfra: Platelet-derived growth factor receptor alpha
Pdgfa: Platelet-derived growth factor alpha
PSR: Picrosirius Red
TAM: Tamoxifen
UC: Ulcerative Colitis
WT: Wildtype

## DATA AVAILABILITY

Normalized transcript count data have been submitted to the National Center for Biotechnology Information Gene Expression Omnibus; GSE252864 and raw sequences to the Sequence Read Archive.

[data/code availability to be added here]

## AUTHOR CONTRIBUTIONS

A.P. designed the research; A.P. and M.P.V conceived research; A.P. drafted edited and revised manuscript; H_o_.W. provided the resources; Z.H. performed experiments, analyzed, and interpreted results of experiments; Z.H., T.H., S.Z., C_r_L., C_i_L., analyzed data; Z.H. prepared figures; A.W., conducted experiments and analyzed data; D.M., S.N., J.B., C.M., K.S., H_a_.W., and K.W. conducted experiments.

---

**Figure S1.**
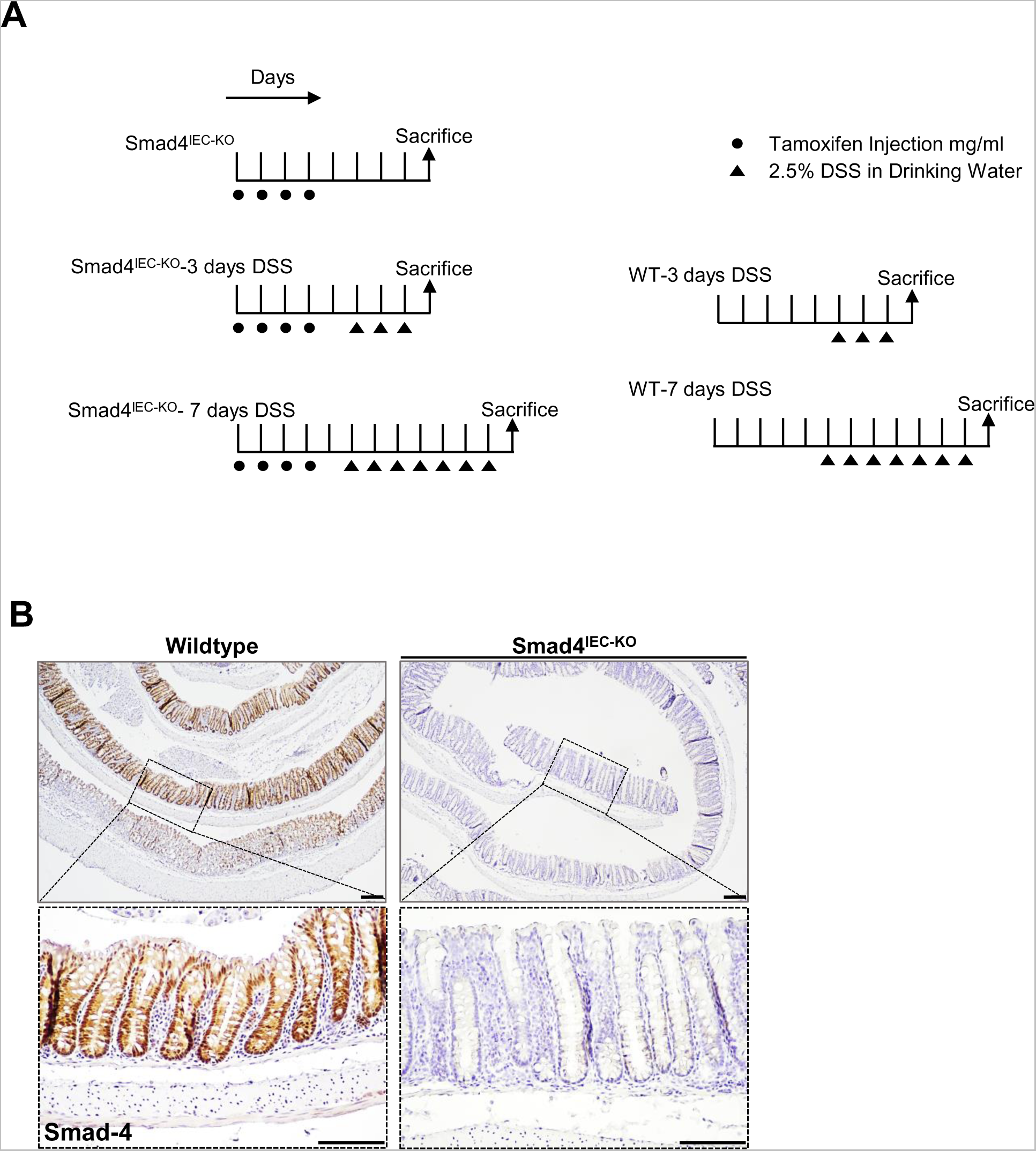
Tamoxifen and/or DSS treatment regimen. (A) Treatment regimen for inducing Smad4 knockout with or without DSS treatment, and DSS treatment in wild type mice are indicated. (B) Smad4 IHC confirming Epithelial- specific loss of Smad4 in the colon following DSS treatment. (n=6: 3 replicates per treatment). Scale bars, 100 μm.

**Figure S2.**
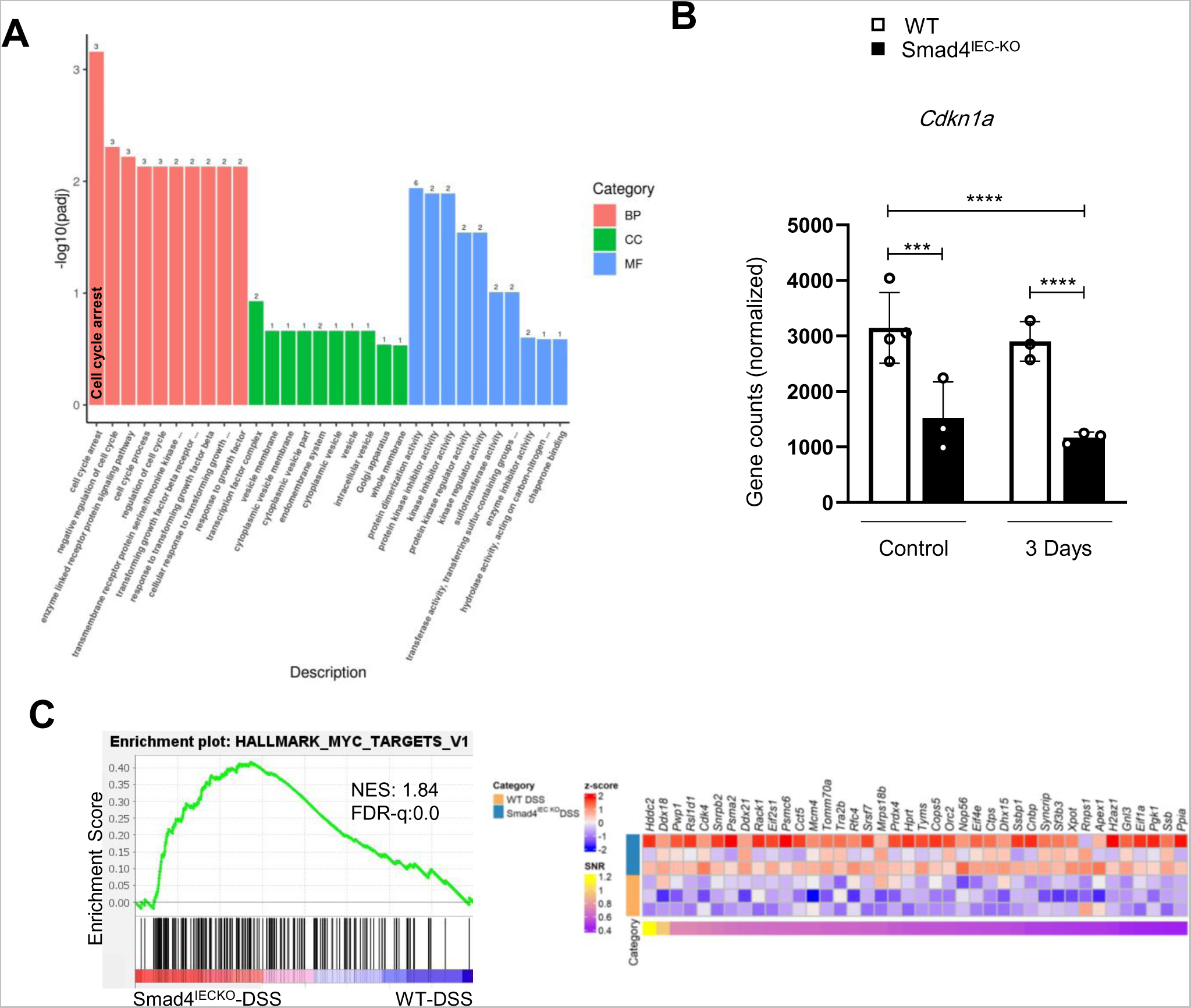
Transcriptomic changes indicating decreased proliferation in the DSS-treated wild type colonic epithelium. (A) Top-enriched Gene Ontology terms among the downregulated gene sets suggesting interrupted proliferation in the DSS-treated wild-type epithelium compared to the DSS-treated Smad4^IECKO^-DSS colon epithelium. (B) Reduced *Cdkn1a* transcript levels in the Smad4^IECKO^-DSS colon epithelium (n=13: 4 replicates per WT and 3 replicates per other treatments); p.adj. values obtained from DEG analysis were used to determine the statistical significance (C) GSEA plot showing enrichment of Myc targets in the DSS-treated Smad4^IECKO^ colon epithelium compared to it wild-type counterpart; the corresponding heatmap shows the core-enriched genes. (n=6: 3 replicates per treatment; *** and **** denote p < 0.001 and p <0.0001, respectively). Scale bars, 100 μm. NES = normalized enrichment score, FDR q= False discovery rate q-value.

**Figure S3.**
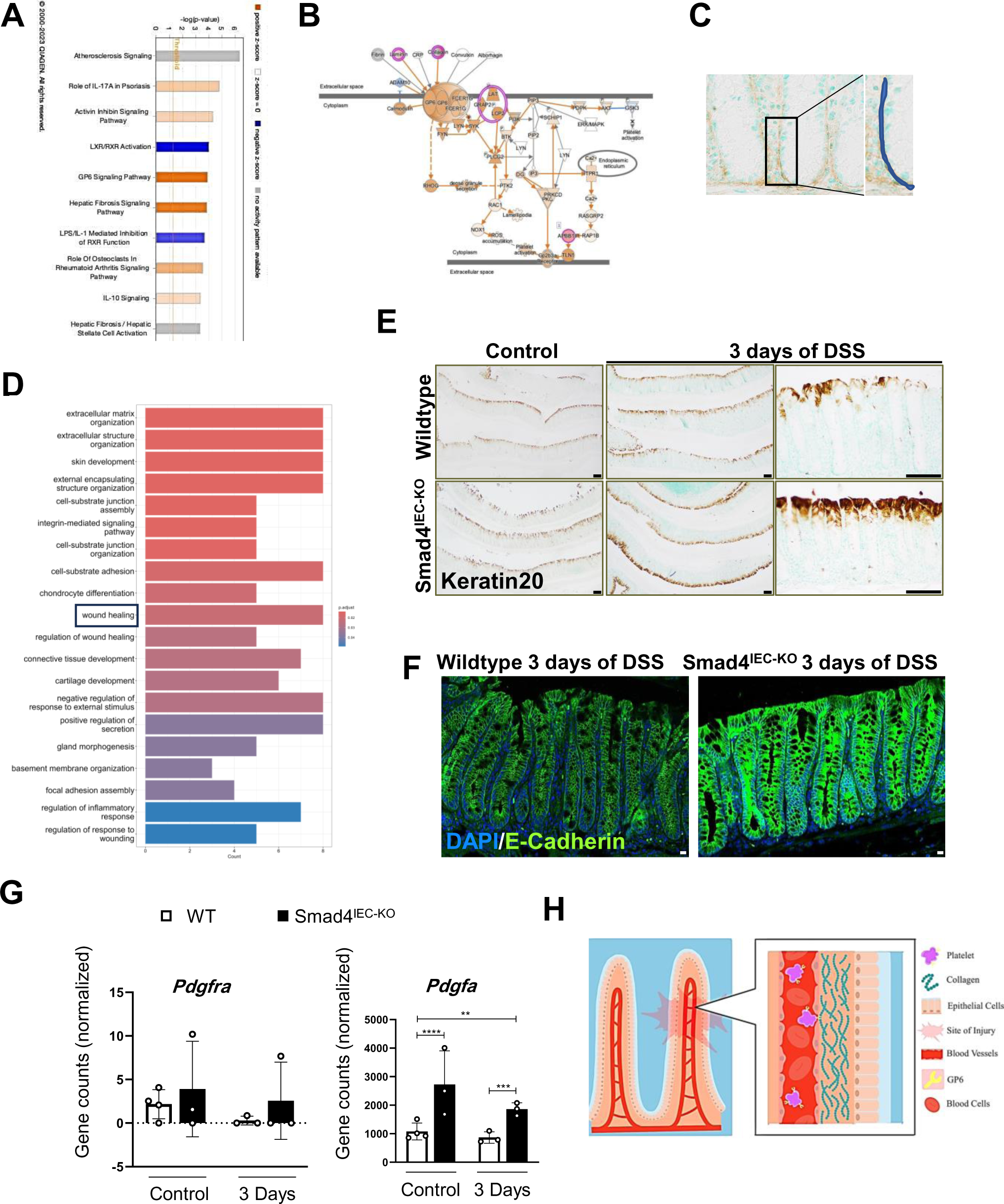
Enrichment of Collagen mediated Gp6 signaling and molecular signatures that support wound healing. (A) Significantly enriched canonical pathways obtained by IPA. (B) Graphical representation of the high-scoring Gp6 signaling pathway as revealed by IPA on the DEGs in DSS-treated Smad4^IECKO^ versus DSS-treated WT epithelium. (C) Mode of quantifying immunoreactivity for Type I collagen in the epithelial ECM. (D) GO terms analysis on the DEGs in DSS-treated Smad4^IEC-KO^ versus DSS-treated WT epithelium showing ECM and wound-healing to be amongst the top-enriched biological processes. (E) Keratin20 immunostaining (n=12: 3 replicates per treatment). Scale bar 100 μm. (F) Confocal fluorescence imaging showing the E-Cadherin (n=6: 3 replicates per treatment). (G) Lack of fibroblast contamination in the epithelial transcriptome assessed by the converse levels of *Pdgfra* and *Pdgfa* transcripts, which encode the *Pdgfra* receptor by fibroblasts and *Pdgfa* ligand by epithelial cells, respectively (n=13: 4 replicates per WT and 3 replicates per other treatments; p.adj. values obtained from DEG analysis were used to determine the significance. (H) Schema illustrating the process of wound healing mediated by GPVI signaling: the interaction between collagen and GP6, subsequently leads to platelet aggregation. |Log2FoldChange| ≥ 0.584 was used to shortlist the DEGs for IPA. Scale bars,10 μm.; **,*** and **** denote p < 0.01, p < 0.001 and p <0.0001respectively)

