## supplementary tables for "Smad4 Loss in the Mouse Intestinal Epithelium Alleviates the Pathological Fibrotic Response to Injury in the Colon"

Table 1. Primary Antibodies used for immunohistochemistry and immunofluorescence staining.

| **Antibody** | **Dilution** | **Catalog#** | **Company** |
| --- | --- | --- | --- |
| α-Smooth Muscle Actin (D4K9N) XP® Rabbit mAb | 1:500 | 19245 | Cell Signaling Technology |
| BrdU | 1:100,1:1000 | M0744 | Dako |
| COL1A1 (E8F4L) XP® Rabbit mAb | 1:200 | 72026S | Cell Signaling Technology |
| E-Cadherin | 1:100 | sc-59778 | Santa Cruz Biotechnology |
| Ki67 | 1:500 | ab16667 | abcam |
| Mouse GPVI Antibody | 1:50 | MAB6758-SP | R&D systems |
| Purified anti-NOS2 | 1:100 | 690902 | Biolegend |

Table 2. Secondary antibodies are used for immunohistochemistry and immunofluorescence staining.

| **Antibody** | **Dilution** | **Catalog#** | **Company** |
| --- | --- | --- | --- |
| Goat anti- mouse IgG (H+L), Biotinylated | 1:300 | BA-9200 | Vector Laboratories |
| Goat anti- rabbit IgG (H+L), Biotinylated | 1:700 | BA-1000 | Vector Laboratories |
| Alexa Flour 488 goat anti-rat IgG (H+L) | 1:250 | A11006 | Invitrogen |
| Goat anti-mouse IgG Alexa Flour 488 | 1:250 | A28175 | Invitrogen |

Table 3. Other reagents and Materials.

| **Name** | **Catalog #** | **Company** |
| --- | --- | --- |
| BrdU (5-Bromo-2'-deoxyuridine) | 19-160 | EMD Millipore |
| DAPI | 5087410001 | Sigma-Aldrich |
| DSS 40 kDa | J63606 | Thermofisher Scientific |
| EDTA | 324504-500ML | EMD Millipore |
| EGTA | 0732-10G | VWR |
| Eosin | 95057-848 | VWR |
| Hematoxylin | 26030-20 | Electron Microscopy Sciences |
| HEPES | J848 | VWR |
| Methyl green | ZH0804 | Vector Laboratories |
| NaCl | S271-1 | Fisher Scientific |
| NaF | A13019-30 | Alfa Aesar |
| Sodium Vanadate | 72060 | Sigma-Aldrich |
| Paraformaldehyde | 15714-S | Fisher Scientific |
| PBS | BP399-4 | Fisher Scientific |
| PMSF | P7626 | Sigma-Aldrich |
| Proteases inhibitor | P8340-5ML | Sigma-Aldrich |
| Tamoxifen | T5648-1G | Sigma-Aldrich |
| Triton x-100 | 0694 | VWR |
| TRIzol | 15-596-018) | Thermo Fisher Scientific |
| 70-micron filter | 431751 | Corning |
| ABC-HRP Vectastain kit | PK-4000 | Vector Laboratories |
| BCA Protein Assay Kit | 0023224 | Fisher Scientific |
| CRP assay kit | ab222511 | abcam |
| Click-iTTM Plus Edu Alexa FlourTM 594 Imaging Kit | 2559157 | Thermo Fisher Scientific |
| ImmPACT (TM) DAB HRP Substrate, | SK-4105 | Vector,Laboratories |
| Sirius red/Fast green kit | 9046 | Chondrex |
